# A high-throughput technique to quantify pathogens’ virulence on the insect model *Galleria mellonella*

**DOI:** 10.1101/297929

**Authors:** Nathalie Parthuisot, Jacques Rouquette, Jean-Baptiste Ferdy

## Abstract

We combined spectrophotometry and an original statistical approach to infer bacteria virulence, using the lepidoptera *Galleria mellonella* as a host model. With this method, it is possible to use a microplate reader to automatize data collection on both pathogens multiplication and host survival on batches of 96 samples.

---

In recent years wax moth, the lepidoptera *Galleria mellonella* (*Pyralidae*), has been widely used for the identification and characterization of microbial virulence factors but also for assessing the efficacy of novel antimicrobial drugs (Tsai et al. 2016; Ramarao et al. 2012; Mukherjee et al. 2013; Mukherjee et al. 2010). It has even been proposed that this animal could provide an increasing alternative to the murine model for studying microbial infections (Goh et al. 2017; Barber et al. 2016). Husbandry of *G. mellonella* is indeed simple, cheap and fast, and its use in research raises less ethical concerns than that of vertebrate models.

In most studies using *G. mellonella*, insect survival is assessed visually, and in some cases with the help of camera and computer assistance. Here we present an alternative to this approach where batches of 96 insects placed in microplates can be followed simultaneously with no human intervention.

*Galleria mellonella* larvae were fed on honey and wax, at 28°C in the dark and in aerated glass jars. In our experiments, we used last instar larvae which were 2-3 cm long and weighted no more than 300mg. Before injection, larvae were starved for 24h and washed with 70% ethanol to limit the risk of co-infection by opportunistic bacteria or fungi.

In the experiments presented here, we used the strain F1D3 of the bacteria *Xenorhabdus nematophila* as a model pathogen (Richards and Goodrich-Blair 2009). This strain expresses GFP (Sicard et al. 2004), which will allow us to quantify bacteria multiplication inside the insect. Insects were injected with 20*µ*l of late exponential LB cultures of *X. nematophila*, with dilution ranging from 10^−^1 to 10^−^3. Six insects injected with sterile culture medium, which were subsequently used as negative controls, were included in each experiment. After injection, each insect was placed in a well of a 96 well glass bottom plate (Sensoplate, Greiner Bio-One). Using glass bottom plates proved to be essential as *G. mellonella* tend to chew holes in plastic plate bottom. Insects were then incubated at 28°C (the optimal growth temperature of *X. nematophila*) for up to five days in a Synergy BioTeK spectrophotometer. At the end of the experiment, the status of each insect (i.e. dead or alive) was visually checked.

We programmed the spectrophotometer so that about every 4 minutes, two signals were measured. GFP fluorescence was measured with excitation set at 485nm and emission at 535nm. Insect autofluorescence was quantified by setting excitation at 330nm and emission at 405nm. We chose this combination of wavelengths because it was weakly affected by bacterial multiplication inside the insect. Autofluorescence signal should therefore be constant over the course of the infection, apart from the noise caused by movements. Exceptions to this rule exist, with some insects displaying a change in autofluorescence as septicaemia developed. But we found this to have no effect on survival estimate.

Insect movements inside a well render fluorescence emissions stochastic. Figure 1A illustrates a typical serie of measurements in control insects that have remained alive for the whole duration of the experiment. Both autofluorescence and GFP fluorescence are highly variable, with no detectable trend during the 72 hours of the experiment shown here. In insects injected with *X. nematophila* (figure 1B), conversely, GFP fluorescence increases sharply as septicaemia develops. Briefly after this increase, stochastic fluctuations cease which indicates that the insect has died.

**Figure 1:**
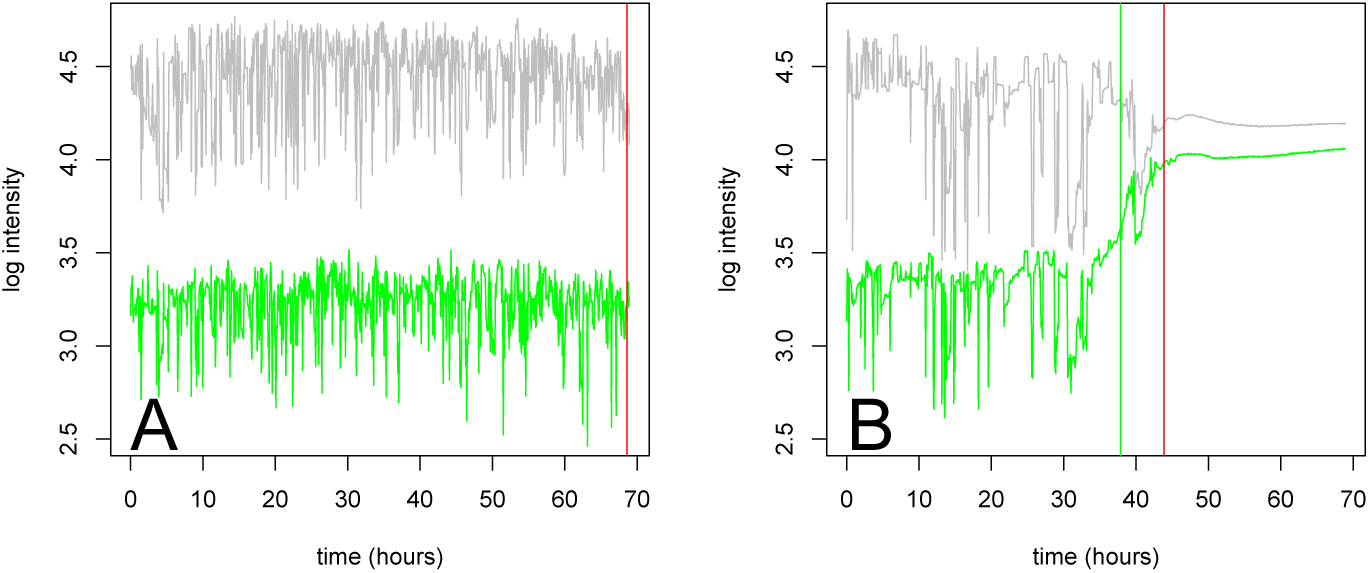
Fluorescence measurements over the 72 hours that follow injection. In panel A, the insect was injected with sterile LB and thus survived. Gray lines indicate autofluorescence while green lines are measurements of GFP fluorescence. Both measurements are highly stochastic, because of insect’s movements inside the microplate. In panel B, the insect was injected with a suspension of *X. nematophila*. After about 35 hours, GFP fluorescence started to increase significantly. After about 44 hours, the variance in both signals dropped to almost zero which indicates that the insect was dead at that time. In both panels, the vertical gray lines indicate the estimated time at which the insect had died. In panel B, the green vertical line indicates the estimated time at which septicemia started.

We propose in the following to estimate time of death as the moment at which variance in autofluorescence signal drops to almost zero. The statistical methods we have developed to do so are implemented in a package, fluoSurv, so that they can be applied using the R software (R Core Team 2018).

Let *Y*_*t*_ be a measure of autofluorescence at time *t* for a given insect; we define *Z*_*t*_ as

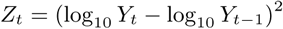

The expected value of *Z*_*t*_ is the variance of log_10_ *Y*_*t*_. We therefore expect a decrease in average *Z*_*t*_ when the insect dies. We fitted a generalized linear model (glm) where *Z*_*t*_ is assumed to be Gamma distributed with average *z*_*b*_ before death and *z*_*a*_ after.

In a first step of the analysis, we estimated *z*_*a*_ as the mean *Z*_*t*_ in the last 5 hours of the experiment for all insects that we know have died during the experiment. We estimated *z*_*b*_ as the mean *Z*_*t*_ over the first 5 hours of the experiment, a time at which we assumed that all insects were alive. Using these two values, we could compute the log-likelihood of the Gamma glm for any time to death *t*_*d*_. We could thus obtain a first estimate 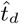 of time to death by maximizing this log-likelihood.

In a second step, we re-estimated *z*_*a*_ as the average *z*_*t*_ for *t* ≥ 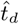 and run again the previous analysis, obtaining therefore a new estimate 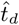. This second estimate (shown as red vertical lines in both panels of figure 1) proved to be more reliable than the first, and was therefore used in all analyses. It should be noted that in most cases the method returns the maximum time value when the insect is alive, as it should be.

Figure 2A illustrates a possible utilization of our method on a complete data set. Here time of death has been computed for each of the 96 insects of a plate. Injecting 96 insects took us about three hours. We thus added the time that elapsed between the first and the last injections to the time recorded by the microplate reader. The estimates of time to death we obtained this way could then be analyzed using, for example, a Cox proportional hazard model. In figure 2, estimates are represented as survival curves, with one curve for each injected dilution.

**Figure 2:**
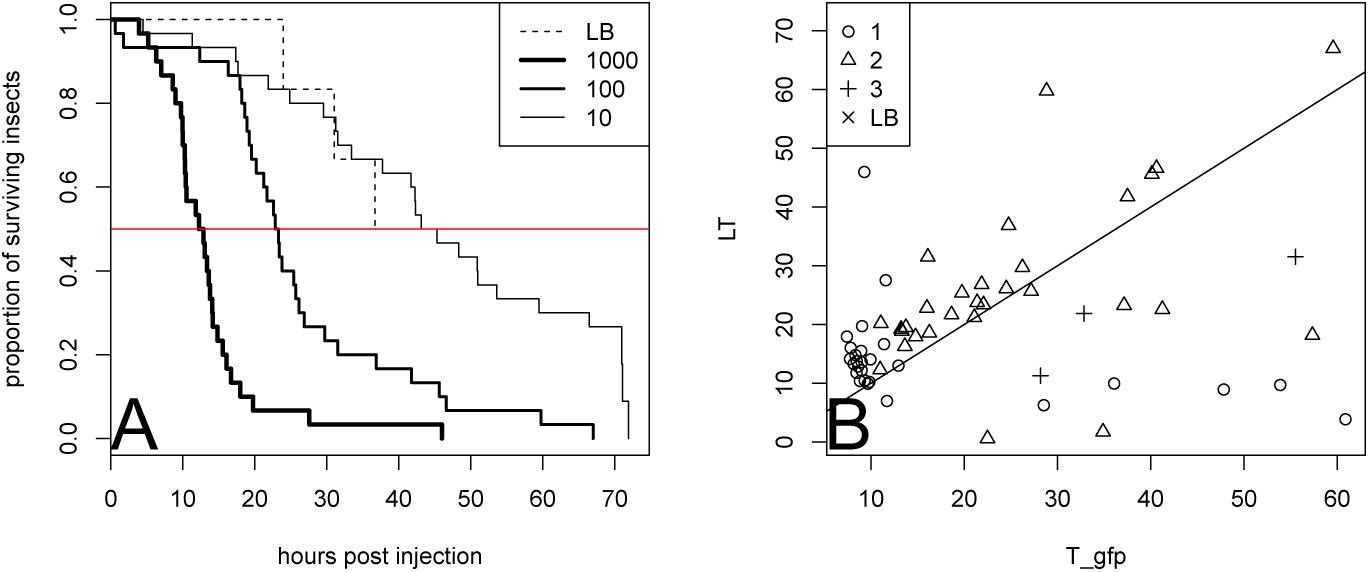
A. Survival curves representing time of death estimates for insects injected with three dilutions of *X. nematophila* cultures. LB corresponds to control insects injected with sterile LB. B. Time to death as a function of the time at which we detected septicemia. This later estimate is obtained by computing when GFP fluorescence exceeds by 10% the maximum intensity value observed over the first five hours of the experiments.

Another use of our method is illustrated in figure 2B, where we represented for each insect both the time to death and the time at which GFP fluorescence started increasing. This analysis demonstrates that in most insects death follows septicemia by a few hours.

The method we describe here allows to estimate time to death simultaneously for large batches of insects using spectrophotometry. From this prospect, it can be considered as a high throughput technique to quantify bacteria virulence using *Galleria mellonella* as a host model. In addition, the use of fluorescent bacterial strains allows to detect bacteria multiplication on the same insects that are used to measure virulence, which offers new experimental possibilities.

